# ERNIE: A Data Platform for Research Assessment

**DOI:** 10.1101/371955

**Authors:** Samet Keserci, Avon Davey, Alexander R. Pico, Dmitriy Korobskiy, George Chacko

**Author notes:** Corresponding author: *Email address:* (George Chacko).

## Abstract

Data mining coupled to network analysis has been successfully used to study relationships between basic discovery and translational applications such as drug development; and to document research collaborations and knowledge flows. Assembling relevant data for such studies in a form that supports analysis presents challenges. We have developed Enhanced Research Network Information Environment (ERNIE), an open source, scalable cloud-based platform that (i) integrates data drawn from public and commercial sources (ii) provides users with analytical workflows that incorporate expert input at critical stages. A modular design enables the addition, deletion, or substitution of data sources. To demonstrate the capabilities of ERNIE, we have conducted case studies that span drug development and pharmacogenetics. In these studies, we analyze data from regulatory documents, bibliographic and patent databases, research grant records, and clinical trials, to document collaborations and identify influential research accomplishments.

## Introduction

Measuring progress in research for reporting, planning, and optimization of research programs is of considerable interest to public and private funders, governments, and research organizations. The broader discussion of measurements of research outputs and research impact has identified challenges and resulted in principles being articulated to guide practice [1, 2, 3, 4, 5]. Quantitative measurements, bibliometrics in particular [6, 7], are frequently used in research assessment to complement and support expert judgment. Beyond citations however, linking to public and commercial data sources such as patents, funding, and regulatory records enables a multidimensional view of the complex research enterprise [8, 9, 10, 11]. Relevantly, a big-data approach involving network analysis of information mined from publicly available and commercial repositories of funding, regulatory, patent, clinical trials, and bibliographic data has been used to document scientific relationships, large collaboration networks, and elite performers contributing to major research breakthroughs such as new therapeutics [12, 13]. In these studies, scientific references in FDA drug approval records, clinical trials, scientific publications, patents, and funding records were used to construct networks of documents connected by citations that enabled descriptions of the large collaboration networks underlying these breakthroughs. This approach identifies influential discoveries as well as high performers in these networks and is extensible to other questions for which a relevant set of documents containing scientific citations exists. Other than relying on subscription services from commercial providers, access to data, particularly research quality bibliometric data [14], presents challenges to large scale data analysis. To support further studies on this theme of citation networks linked to other data sources, we have developed Enhanced Research Network Informatics Environment (ERNIE), a simple, extensible, cloud-based platform that leverages modern data science and open source software to aggregate data at scale for discovery. ERNIE contains FDA records, clinical trials and clinical guidelines data, bibliographic and patent data, and funding records from the National Institutes of Health. A modular design enables the addition, deletion, or substitution of free or leased data sources by users. The platform consists of a database and associated methods for generating and analyzing networks constructed from its data. We illustrate its utility with two examples i) through reproducing prior studies [12] (ii) tracing the evolution of a pharmacogenomic diagnostic kit [15] from its origins in microarray technology [16, 17, 18].

## Materials and Methods

### Infrastructure

ERNIE is hosted in the Microsoft Azure cloud. The primary server, a standard D8s v3 (8 vCPUs, 32 GB RAM) with 8 TB of attached SSD storage, acts as a central hub for data related processes and hosts data in a PostgreSQL 9.6 relational database. Secondary computing support is provided through (i) a standard DS4 (8 vCPUs, 28 GB), CentOS 7.4 with Apache Solr 7.1 installed that is used to index and search content (ii) a standard D16s v3 (16 vCPUs, 64 GB memory), CentOS 7.4 with the Neo4j graph database installed for graph analytics (iii) an Apache Spark cluster with 9 nodes that is provisioned on demand using HDInsight and is used for compute intensive tasks with workflows that use Apache Sqoop, and PySpark. Access is secured through SSH tunneling and two-factor authentication. Interactive users have access to Python, SQL, R, Java, Jupyter notebook, and standard Linux utilities.

### Data sources

Data in ERNIE are derived from both publicly available and commercial sources and are stored in a documented relational schema. Publicly available data are copied from ClinicalTrials.gov [19], the US Food and Drug Administration (FDA) Orange [20] and Purple [21] Books, the United States Patent Office [22], NIH ExPORTER [23], and the National Guideline Clearinghouse [24]. Leased data were acquired from Clarivate Analytics and consist of the Web of Science Core Collection [25] and the Derwent World Patent Index [26]. Custom Extract-Transform-Load (ETL) scripts developed in Bash, Python, Java and SQL are used for initial loading as well as refreshes. Web of Science, Patents, Clinical Trials, NIH ExPORTER, and Clinical Guidelines data are updated weekly. All other data sources are updated monthly. Automated processes are managed by the Jenkins Continuous Integration server, which is used to (i) deploy scripts dedicated to data transformation (ii) direct construction of Solr indexes as new data is introduced into ERNIE (iii) build citation networks for case studies (iv) manage data refreshes. Source code and procedures are available under the open-source MIT license and are available on GitHub [27]. Table 1 provides an overview of records in the major data sources (excluding records from tables used to process data) in ERNIE.

**Table 1:** Summed counts of records in ERNIE grouped by major data sources. The number of records for each data source is expressed in scientific notation with coefficients rounded to one decimal place. The count of records includes related records within the schema. For example, the Web of Science Core Collection consists of approximately 6.7 x10^7^ publication records with associated records in related tables that include author names, cited references, and synonymic identifiers that add up to the total of 2.5 x10^9^ shown above.

|  | Data Source | No of Records |
| --- | --- | --- |
| 1 | Web Of Science Core Collection | $2.5 \times 10^9$ |
| 2 | Derwent US Patents | $2.4 \times 10^8$ |
| 3 | NIH ExPORTER | $7.1 \times 10^6$ |
| 4 | NLM Clinical Trials | $8.6 \times 10^6$ |
| 5 | FDA | $5.0 \times 10^4$ |
| 6 | AHRQ Clinical Guidelines | $2.6 \times 10^3$ |

### Data architecture and governance

ERNIE data is drawn from several upstream sources of varying quality. Cleansing approaches in place are designed to favor data quality and normalized data over completeness. Key aspects of ERNIE’s data architecture and governance are (i) De-duplication to eliminate logically duplicate records, that is records with identical primary or unique keys. In ERNIE, duplicate records from such a group are deleted while leaving in place a single representative record (preferring a record which is updated last) (ii) Normalization to establish foreign key relationships between children and parent entities and eliminate orphans: child records with non-existing parents. (iii) Schema permanence to improve availability of data and simplify data schema management (iv) Data stewardship to assign responsibility for quality of a particular data subject area. Data stewards are notified when a data issue is detected and drive the workflows necessary for data issue remediation. They are also typically responsible for communicating changes in data structures.

### Workflows

The core analytical workflow in ERNIE can be divided into five stages (i) Source document identification, in which a set of source documents relevant to the question being asked is assembled, e.g., FDA regulatory reports or policy documents (ii) Citation extraction and matching in which text references are extracted from the source documents in the first step and matched to unique identifiers such as PubMed ID (pmid), UT (Web of Science Accession Number), or US patent numbers. (iii) Amplification and linking, in which publications and patents linked by citation to extracted references are identified through citation records in the Web of Science and linked to other records in ERNIE. (iv) Network construction, in which the data assembled through the preceding stages are represented as a network. (v) Analysis for discovery and visualization of the network data to stimulate further investigation.

### Source document identification

A set of source documents relevant to the question being asked is assembled and searched for citations to scientific literature. The process involves expert judgment, is subjective, and centers around regulatory, policy, and technical documents. For example, FDA documents related to approvals of the therapeutics ipilimumab and ivacaftor were mined for scientific references.

### Citation extraction and matching

References to scientific literature and patents found in the source documents are matched to unique identifiers using a manual search of PubMed or partially-automated search of the Web of Science Core Collection. The product of reference extraction and citation matching is termed the ‘seed set’. Manual extraction involves searching for text strings that represent references of interest and matching those to unique identifiers in PubMed and/or Web of Science. For example, US6984720 B1, the core patent relevant to ipilimumab contained 52 references to scientific publications listed under non-patent citations. All 52 of them were manually matched to PubMed identifiers (pmid) using the PubMed website. In a partially-automated approach, Apache Solr is used to to create and store indexes of ERNIE data that can facilitate user searches that return informative, ranked, and scored query results and is the basis for a process that takes advantage of indexed Web of Science publications in ERNIE. A similar approach has been described for citation matching of raw text strings [28]. In our process, a text file containing references is passed as input to a Python script. The user manually selects matching publications from the output hit list that includes citation information, an UT, and the numeric output of the Solr scoring function. In the case of ivacaftor, US7495103, its core patent contained 20 references to scientific publications. Using the manual PubMed approach, 8 of 20 ivacaftor citations were correctly identified. Using the Solr indexing approach, 15 of 20 references were correctly identified. In an extension of this process, selection is also made using a percentile approach, e.g. nth percentile of Solr scores for a given dataset calibrated by scores for known true positive and true negatives. The partially-automated approach process is more scalable and faster than the manual one. A combination of these approaches was used in the case studies described.

### Amplification

Once unique identifiers to scientific literature are assembled in the seed set, cited and/or citing references in the Web of Science Core Collection are identified using the relational data in ERNIE. Where possible, UTs are mapped to pmids using a lookup table and then to grants using data from NIH ExPORTER. References are then linked to authors using another table in our schema. The author address field in Web of Science is used to identify author countries, which were then standardized by using a string matching approach to match to a lookup table constructed from a current list of countries published by the US Department of State. Cases where a Levenshtein distance of greater than 1 between country name in Web of Science data and the lookup table were managed through ad-hoc scripted corrections, e.g., ‘FED REP GER’ was converted to ‘GERMANY’.

### Workflows and Network analysis

For the ipilimumab and ivacaftor case studies, a single generation of cited references was extracted from an input of pmids and/or UTs. The seed set was derived from mining FDA regulatory documents, clinical trials data, patent literature, and bibliographic databases for references that approximately preceded the date of FDA approval for these two drugs. These references were then combined into a citation network where nodes are publications and authors. Grant support, and institutional location are attributes of these nodes. A single verb, ‘cites’, was used to draw edges between nodes, e.g., node A cites node B. An optional step, availed of in these two cases, was to include the cited references in review articles published within a year of the FDA approval for these drugs.

For network analysis in the first two case studies on drugs, four input files are used that are derived from the amplification stage (i) article and author edge list, a mapping of publications to authors (ii) a single generation of cited or citing references derived from the amplification cycle (iii) publication and publication edge list, the citation network after amplification stored as source-target citation pairs (iv) publication and published year, a mapping of publications to their year of publication.

Once these four files are generated, the network analyzer code generates output files (i) article scores based on weighted citation counts from within the network. (ii) a list of author scores based on summed article scores. (iii) an edge list of articles for visualization. (iv) descriptive statistics, this file contains descriptive statistics about articles and authors generated during network analysis. The process is automated and managed through a Jenkins workflow. As we have previously noted [13], the use of pmids for network construction results in significant loss of information, thus we used Web of Science data for network construction. For each article, we calculated in-network citations, as well as weighted citations (sum of article citations plus sum of citations of an article’s first degree neighbors). For each author, we calculated the number of articles, total citations and propagated in-degree rank (PIR) [12].

For the pharmacogenomics case study, we mapped citation paths from the DMET Plus diagnostic panel for drug metabolism to a set of references that define the origins of the DNA microarray technology. We incorporated a combination of references cited b the DMET Plus set and references citing microarray set then generated edgelists and nodelists directly in PostgreSQL and exported the data to the Neo4j graph environment. These data were then analyzed with Cypher queries.

### Visualizations

Visualizations were performed using Cytoscape [29] using edgelists and nodelists as input. Edgelists were generated from graph network data as source-target node pairs with additional columns source type (stype) and target type (ttype) that qualify nodes. Node types used in these studies were root, publication (wos id), clinical trial (ct), fda document(fda), author, and patent. An example of a source target pair is provided below that represents the citation of a publication in a patent.

| source | stype | target | ttype |
| --- | --- | --- | --- |
| US7824894 | patent | WOS:A1996UX51900025 | wos_id |

## 1. Results and Discussion

### Ipilimumab and ivacaftor

To establish a baseline for content and to validate our workflow, the FDA-approved therapeutics ipilimumab and ivacaftor were selected as case studies since they have previously been studied [12]. Williams and colleagues reported that ‘the citation network leading to ipilimumab includes 7,067 different scientists who listed 5,666 different institutional and departmental affiliations and includes discoveries spanning 104 years of research. Results for ivacaftor were similar: 2,857 different scientists from 2,516 different institutional and departmental affiliations, with discoveries spanning 59 years of research.’ Using data in ERNIE and our improved workflows, we report findings on a larger scale, 56,530 publications for ipilimumab representing the contributions of 122,770 scientists from 105 countries that span 110 years of research and 9,292 publications for ivacaftor representing the contributions of 33,320 scientists from 93 countries that span 105 years of research (Table 2). The majority of the publications in both these case studies correspond to the period 1970-2010.

**Table 2:** Ipilimumab and ivacaftor. The count of records for publications, authors, years, and countries for the ipilimumab and ivacaftor case studies is compared with historical data from the study of Williams (2015),

| Case Study | Publications | Authors | Years | Countries |
| --- | --- | --- | --- | --- |
| ipilimumab (this study) | 56,530 | 122,770 | 110 | 105 |
| ipilimumab (2015) | 1,919 | 7,067 | 104 | - |
| ivacaftor (this study) | 9,292 | 33,320 | 105 | 93 |
| ivacaftor (2015) | 934 | 2,857 | 59 | - |

Although we recorded many more researchers in these two networks, the list of researchers generated by Williams overlaps heavily with the list of researchers in our study. For the Top 20 researchers among elite performers in the Williams study as determined by PIR scores, all 20 for ipilimumab and 19 for ivacaftor are found in our list of researchers. However, the two lists are not identically ordered. For example, 50% of the top 10 researchers in our ipilimumab set overlap with the top 10 in the Williams dataset. We attribute these differences to improved harvesting of references and richer content of the data in ERNIE and suggest that the use of bins is likely to be more useful than individual rank order comparisons in identifying elite performers.

We observed instances of missing author address data from the publications in our case studies particularly for years around 1950-1970 and this trend is also observed in the 67 million publications in the Web of Science dataset that we have in ERNIE (Fig. 2 Right Panel). While these data are missing from the Web of Science Core Collection, the effect of the missing values is is likely to be significant when analyzing institutional and city affiliations in the 1950-1970 period rather than the 1970-2010 period from which most of the references in these two datasets are drawn.

**Figure 1:**
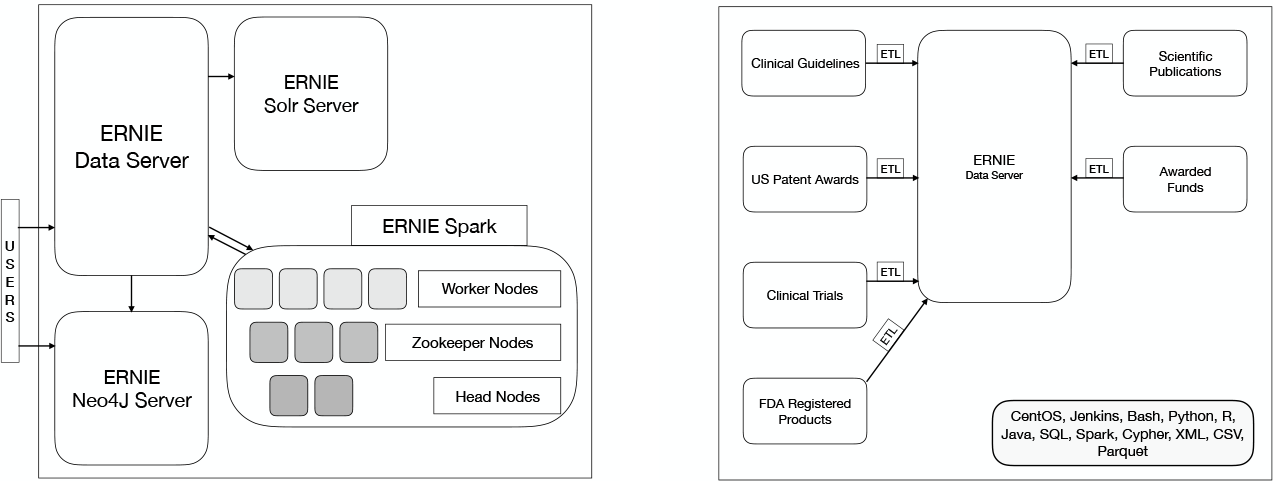
System Context View. ERNIE runs on a PostgreSQL 9.6 database in a CentOS 7.4 environment in the Microsoft Azure cloud. Users access the data server or the Neo4j server through through multifactor authentication (MFA) *Left Panel* Support for text searching is provided through an Apache Solr server. Graph analytics are conducted on datasets exported from PostgreSQL to the Neo4j server. An Apache Spark cluster is provisioned for compute intensive tasks. *Right Panel* ERNIE data is populated through automated custom ETL processes developed in Python, Bash, Java, and SQL for data in XML or CSV

**Figure 2:**
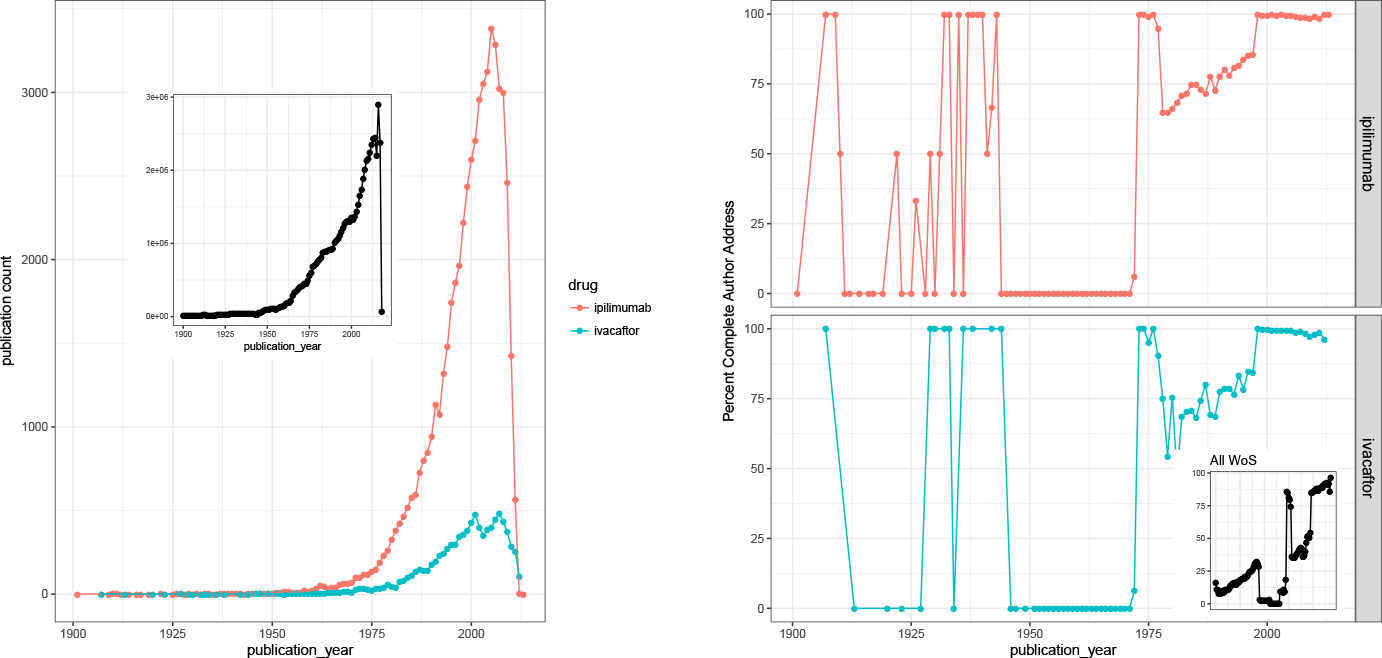
Features of ipilimumab and ivacaftor networks. Content and workflows in ERNIE were validated by reproducing prior work on the FDA approved drugs ipilimumab and ivacaftor. *Left Panel.* Counts of publications in each network are shown by year of publication *Inset* Counts of publications for the Web of Science Core Collection are shown. *Right Panel.* The percentage of publications in each network that have author addresses is shown for each year. *Inset* The percentage of publications with author addresses is shown for each year for approximately 67 million publications in the Web of Science Core Collection.

To identify influential publications, we calculated a weighted citation score for each node in these networks [12, 13], which consists of in-network citations to each node summed with in-network citations of its first degree neighbors. These data are shown in Figure 3 as maximum, minimum, and average values of all publications in a given year.

**Figure 3:**
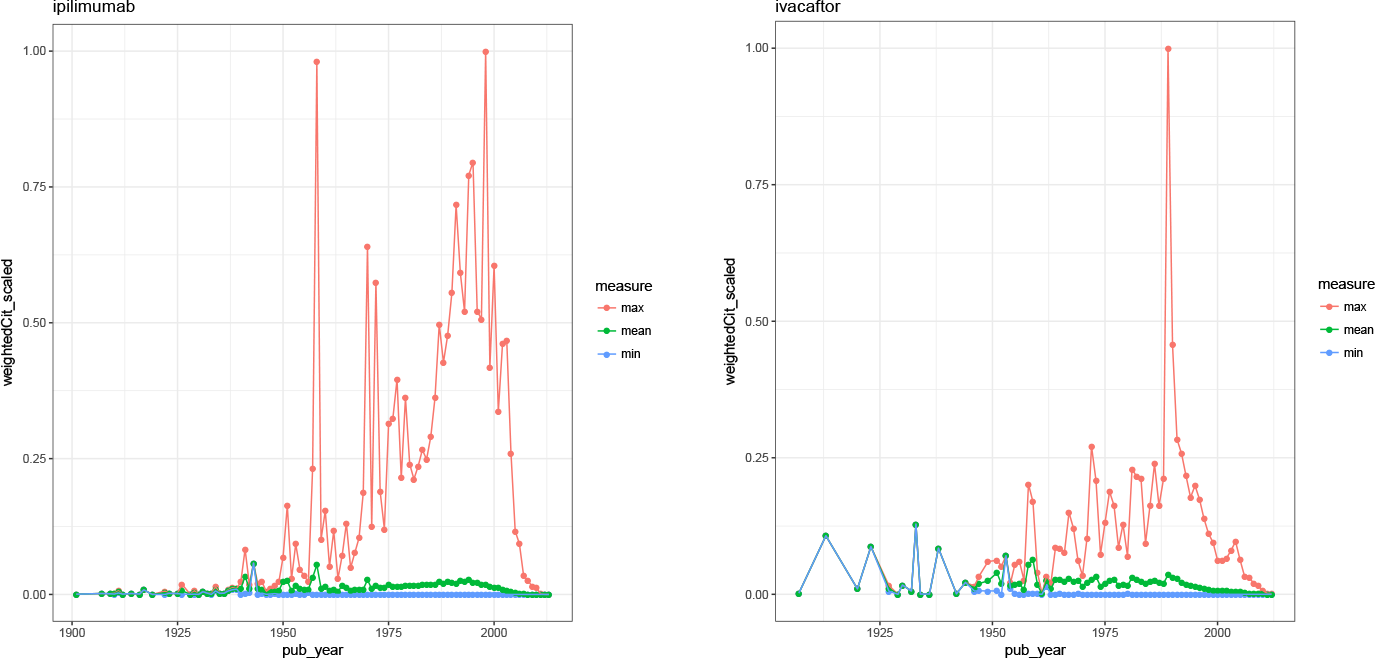
Influential Publications. Content and workflows in ERNIE were validated by reproducing prior work on the FDA approved drugs ipilimumab (Left Panel) and ivacaftor (Right Panel). Weighted citations were were calculated for publication in each network as the sum of citations for each publication and its first degree neighbors then scaled between 0 and 1 for comparison across networks. Maximum, minimum, and mean weighted citations for publications are grouped by year of publication

In a variant of the approach used to study ipilimumab and ivacaftor, we also conducted case studies on buprenorphine and naltrexone, drugs used in the treatment of substance abuse (Supplementary material). For these two case studies, the seed sets were assembled by a simple keyword search in PubMed followed by extraction of a generation of cited references. The resultant network was analyzed for acknowledgment of grant support from NIH.

### Citation Pathways

Microarray technology has had considerable impact in biomedical research. A keyword search for ‘microarray’ in 2018 using PubMed yields more than 86,000 references, testifying to its remarkable impact that has been documented in historiographic studies from various perspectives [30, 31, 32]. The technology has since been applied to pharmacogenomics, and diagnostic kits are now available to used to predict phenotypes in drug metabolism [33] based on polymorphisms in cytochrome P450 genes.

To study the evolution of this application of microarray technology, we assembled a dataset in the PostgreSQL environment and exported it to a graph database environment to facilitate analysis. We constructed traces of citations between nine publications representing the origin of the technology [16, 17, 18] dating from 1991 and earlier, to 28 references defining the more recently developed DMET Plus [15] diagnostic kit. From the microarray references, two successive generations of citing references were extracted using Web of Science data. From the diagnostic references, two successive generations of references were extracted. The data were combined and exported into the Neo4j graph database to create a graph with 394,544 documents represented as nodes.

The PageRank algorithm has been used to identify influential publications in citation networks including those described as ‘gems’ [34] and has the theoretical advantage of considering all nodes in a network when scoring as compared to our weighted citation approach that considers only first degree neighbors. Thus, each node was annotated with PageRank scores, as well as acknowledgement of grant support from NIH institutes. Paths were traced from all nodes in the microarray set to all nodes in the diagnostic set using the allShortestPaths function in Neo4j. Additional constraints were also applied to count shortest paths that traversed at least one node in the 90th percentile of PageRank scores for this distribution and/or at least one node acknowledging NIH support. These enabled identification of paths that passed through at least one highly influential node that included support from NIH.

The results are summarized in Table 3 and indicate that of 2,434 shortest paths, roughly 68% of the paths from microarray technology to DMET Plus references pass through at least one node funded by NIH; 64% pass through a node with a PageRank score in the 90th percentile; and 46% of the paths pass through at least one node with a PageRank score in the 90th percentile *and* at least one node funded by NIH implying a significant role in supporting the evolution of microarray technology to a screening for cytochrome P450 polymorphisms.

**Table 3:**
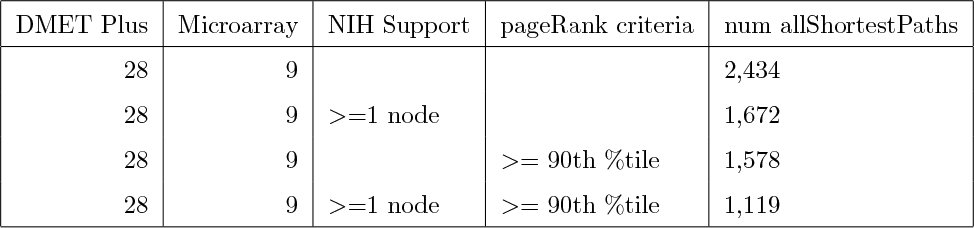
Citation pathways between DNA microarrays and DMET Plus panel for pharmacogenomic typing. The number of pathways traced using the allShortestPath function in Neo4j is shown between a set of 28 publications that describe the Affymetrix DMET Plus pharmacogenomic diagnostic panel. Additional constraints were applied by requiring each path to pass through at least one NIDA or other NIH supported node and/or a node with a PageRank score in the 90th percentile or greater.

| DMET Plus | Microarray | NIH Support | pageRank criteria | num allShortestPaths |
| --- | --- | --- | --- | --- |
| 28 | 9 |  |  | 2,434 |
| 28 | 9 | >=1 node |  | 1,672 |
| 28 | 9 |  | >= 90th %tile | 1,578 |
| 28 | 9 | >=1 node | >= 90th %tile | 1,119 |

To focus this analysis further towards DNA microarrays, we also counted shortest paths between a single publication demonstrating the use of oligonucleotide arrays for rapid DNA sequence analysis [18] and the DMET set that also indicates a significant role for NIH supported research in the knowledge flow between microarray technology and a specialized application of it and observed that 57% of the 258 shortest paths contained at least one node with NIH support (Figure4).

**Figure 4:**
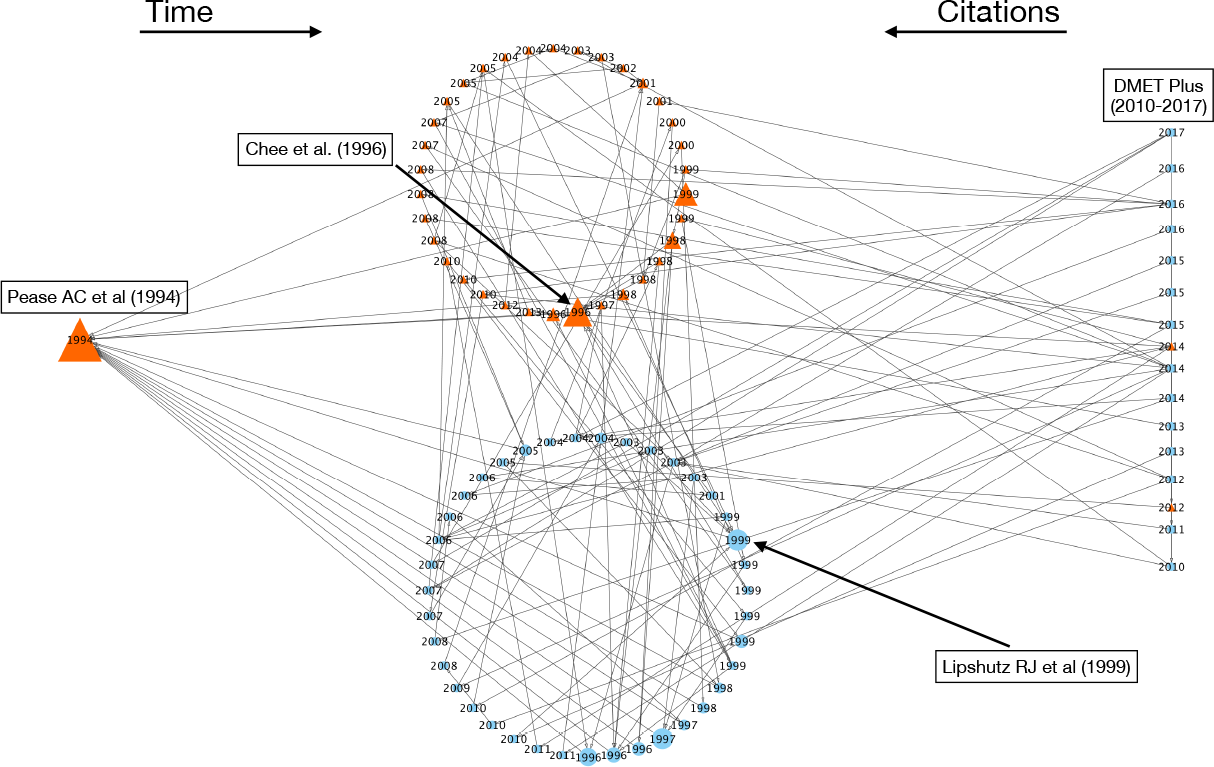
Tracing Paths of Knowledge Transfer from Technology Application to Origin. The allShortestPaths function in Neo4j was used to trace paths from a set of 28 publications published between 2009 and 2018 that describe the DMET Plus pharmacogenomic typing panel to a single publication that describes sequence analysis using light-synthesized dense oligonucleotide arrays in 1994. 258 shortest paths were detected in a graph of 394,544 nodes and 676,591 relations of which 153 paths (57%) passed through at least one publication that acknowledged support from a US Dept of Health and Human services agency including the National Institutes of Health and 50 of these paths (18%) passed through at least one node in the 90th percentile of the distribution of PageRank scores *and* at least one node acknowledging NIH support. Nodes are sized according to PageRank scores and orange colored triangles indicate US DHHS support. Examples of nodes with high PageRank scores are labeled.

## Conclusions

The availability of affordable computing, data science methods, and data exerts a democratizing effect on research activities [35]. Building upon prior art, we have sought to demonstrate the relative ease with which multiple sources of publicly available and commercial data can be integrated to enable studies of scientific advances through the assembly of document sets that are linked by citations. We have demonstrated two variations on a simple theme of data mining, network representation, and analysis. Importantly, the analytical process supports expert judgment. The ERNIE paradigm is intended to support distributed access, curation, and ownership of evaluation datasets. Smaller evaluation teams unable to afford commercial services may be able to work with subsets of the content in ERNIE, e.g., PubMed alone instead of PubMed and Web of Science and modify the code and workflows to their own needs. We are exploring other linking approaches based on text similarity.

We also provide initial observations in these case studies that can be extended through new analytical approaches and additional data sources. We do not claim that these studies are complete. One limitation is incomplete data on funding; we focused on NIH because of its very large spending on biomedical research and the availability of its funding records but significant spending by other funders is not accounted for. This limitation could be addressed in part by improved extraction from Web of Science data, which contains information on other funders. An alternative is subscription to a commercially available source of information on funding. A second limitation is that the results from our case studies are largely descriptive, which may be appropriate for an article focused on demonstrating features of a data repository. However, it is also important to rigorously analyze these datasets further and we have ongoing efforts in this regard. A third caveat is the focus on citations, which have their own limitations [2, 7]. Our case studies themselves would be more informative if conducted in close collaboration with the policy community to help refine the questions to be asked and interpret the results using expert judgment.

## Acknowledgments

Samet Keserci’s contributions to this manuscript were made when he was a full time employee of NET ESolutions Corporation. Research and development reported in this publication was partially supported by Federal funds from the National Institute on Drug Abuse, National Institutes of Health, US Department of Health and Human Services, under Contract No. HHSN271201700053C. The content of this publication is solely the responsibility of the authors and does not necessarily represent the official views of the National Institutes of Health. Clarivate Analytics Inc, the vendor of Web of Science and Derwent World Patents Index had no role in the funding, experimental design, review of results, and conclusions presented in this paper. We also thank two anonymous reviewers who reviewed an earlier version of this manuscript for for their constructive criticism.

